# Method-dependent biases in cell type detection between single-cell and single-nucleus RNA sequencing in the photosymbiotic acoel *Praesagittifera naikaiensis*

**DOI:** 10.64898/2026.09.27.754747

**Authors:** Ryo Nakamura, Mayuko Hamada, Kenji Kobayashi, Satoshi Ansai, Mirco Dindo, Tosuke Sakagami, Yi-Jyun Luo, Yutaka Satou, Toshio Sekiguchi, Tatsuya Sakamoto

## Abstract

**Background:** Comparisons of single-cell and single-nucleus RNA sequencing (scRNA-seq and snRNA-seq) data have been described in some mammalian tissues and, subsequently, in *Drosophila*, but remain unexplored in most invertebrate lineages. The xenacoelomorphs occupy key phylogenetic positions, yet they differ anatomically from mammals. They have a reduced extracellular matrix, high-salt body fluid, and no circulatory system. Despite these differences, they possess a well-developed nervous system. One of the xenacoelomorphs, the photosymbiotic acoel (*Praesagittifera naikaiensis*) also harbours symbiotic *Tetraselmis* algae, whose RNA can be co-captured with host RNA.

**Results:** We compared scRNA-seq and snRNA-seq data from whole *P. naikaiensis* specimens. Both methods yielded high-quality data with comparable gene detection but a larger share of scRNA-seq reads derived from symbiotic algae. Gene-level analyses revealed that neural genes were enriched in snRNA-seq relative to non-neural genes. Cross-method label transfer and integration-based validation identified six snRNA-seq clusters, including some neural populations, that lacked a clear scRNA-seq counterpart. In contrast, three cell populations, including muscle and metabolically active clusters, were reciprocally validated as captured by both methods.

**Conclusions:** Our results show that key snRNA-seq advantages, particularly the enhanced recovery of neural transcripts, are recapitulated in our dataset, which is consistent with previous reports in mammals. Furthermore, snRNA-seq reduces symbiont-derived reads and recovers several cell populations underrepresented in scRNA-seq. These findings provide practical guidance for cell atlas construction in non-model, symbiotic invertebrates.

## Background

Single-cell and single-nucleus RNA sequencing (scRNA-seq and snRNA-seq) have become important tools for dissecting cellular heterogeneity. Systematic comparisons have shown method-dependent biases in mammalian systems [1–7]. Specifically, snRNA-seq excels for profiling neurons from brain tissue [8–10] and hard-to-dissociate tissues (e.g., kidney [6]), whereas scRNA-seq can capture cytoplasmic transcripts [1]. Among invertebrates, such direct comparisons remain limited to *Drosophila* [11, 12], leaving critical gaps for other invertebrate lineages, especially for early-branching groups that occupy key phylogenetic positions and exhibit highly different body organisation.

Xenacoelomorpha, a clade occupying a contentious but phylogenetically informative position, proposed either as sister to all other Bilateria [13, 14] or to Ambulacraria [15–18] substantially differ from mammalian systems in tissue architecture. They exhibit reduced or lacking extracellular matrix (ECM) including in the nervous system, and they lack circulatory and vascular systems [13, 19, 20]. Despite this simplicity, they possess developed and centralised nervous systems [20–27]. Whether the method-specific biases in mammalian tissues apply to such organisms with anatomically different body plans remains unknown. Understanding method performance in these lineages is essential not only for constructing accurate cellular atlases but also for enabling robust cross-species comparisons to trace the evolution of cell types and tissue organisation across Metazoa.

As a member of Xenacoelomopha, the marine acoel (*Praesagittifera naikaiensis*) has a simple body plan with a centralised nervous system [27] and its small size enables whole-animal profiling. In addition, *P. naikaiensis* harbors photosymbiotic algae (*Tetraselmis*) (Fig. 1A) [28], a feature also found in other acoels such as *Symsagittifera roscoffensis* [21, 22], allowing us to evaluate the impact of non-host RNA contamination on each method.

**Figure 1.**
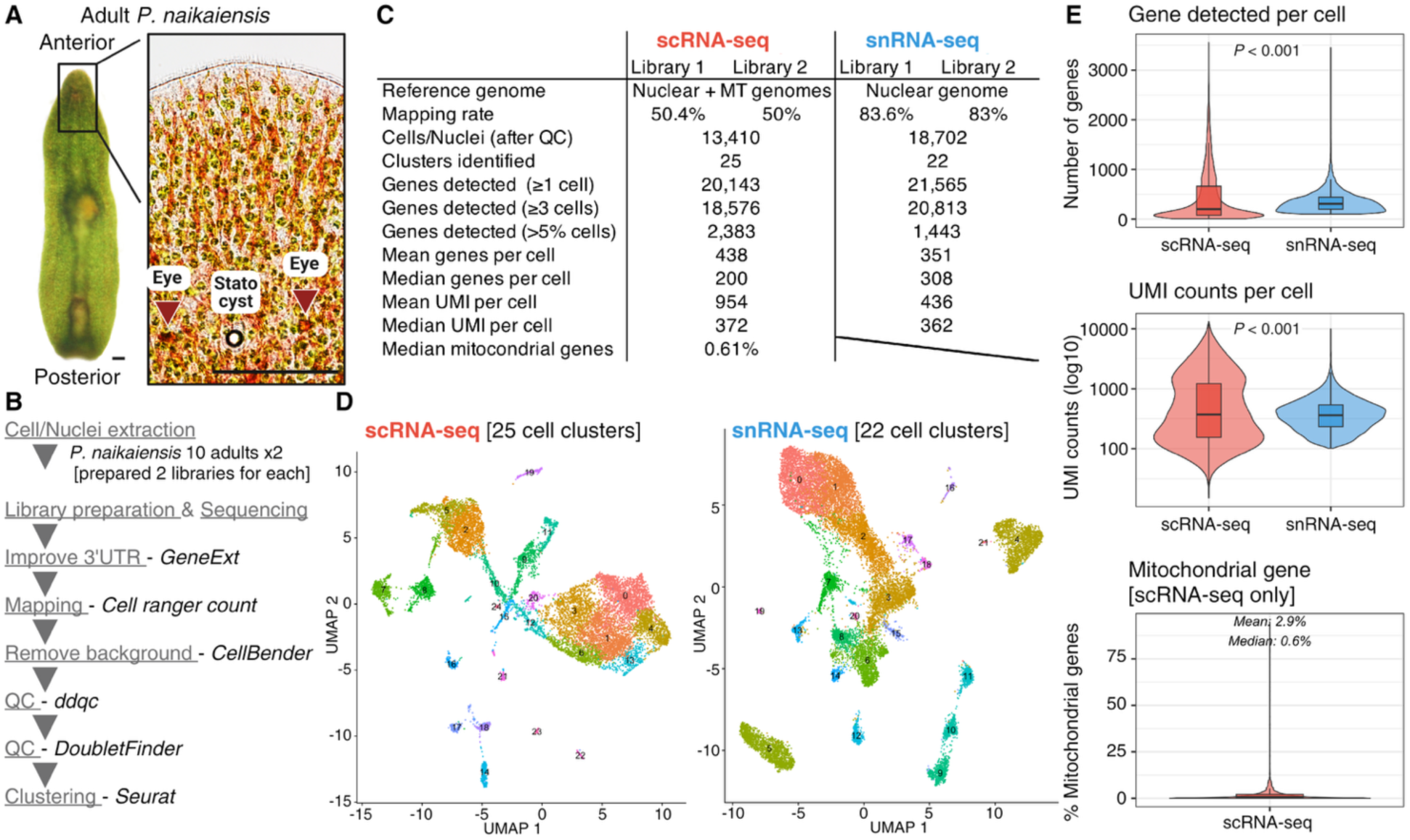
Dataset overview and quality metrics of scRNA-seq and snRNA-seq in a xenacoelomorphan adult *P. naikaiensis*. **(A)** *P. naikaiensis* adult. The anterior region (right panel) contains the sensory organs (statocyst and eyes) that respond to gravity and light, respectively. All green dots show symbiotic algae. Scale bars, 100 μm. **(B)** Workflow for scRNA-seq and snRNA-seq datasets. **(C)** Summary statistics along data analyses. Note the substantially different mapping rates (scRNA: ∼50%; snRNA: ∼83%), reflecting symbiotic algal RNA content in whole-cell preparations. Both methods detected >20,000 genes with high data quality. **(D)** UMAPs of scRNA-seq (left) and snRNA-seq (right), generated by Seurat. **(E)** Quality control metrics from both methods. Violin plots showing distributions of host genes detected per cell/nucleus (top), UMI counts (middle), and mitochondrial gene percentage (bottom, scRNA-seq only). scRNA-seq shows greater heterogeneity in gene detection.

Here, we present the comparison of scRNA-seq and snRNA-seq not only in non-model invertebrates but also in photosymbiotic animals. We profiled whole *P. naikaiensis* specimens using both methods to (1) identify gene categories and cell types preferentially detected by each method and (2) assess method-dependent differences and cross-method consistency, including the effects of co-captured algal RNA. Neural genes and several cell populations differed between methods, whereas others were captured by both, providing a reference for planning cell-atlas studies in invertebrates.

## Methods

### Cell extraction and fixation for scRNA-seq

Adult *P. naikaiensis* individuals were collected and processed for scRNA-seq using a methanol-acetic acid fixation protocol. Briefly, 50 worms were collected in 1.5 mL low-binding tubes and washed twice with 500 µL artificial seawater (ASW). Tissues were dissociated by incubation with 200 µL of 0.1% trypsin (Sigma-Aldrich #T4799) in calcium- and magnesium-free seawater (CMF) for 10 minutes at room temperature with gentle inversion. Enzymatic digestion was quenched by addition of 250 µL CMF containing 1% BSA (final concentration; Sigma-Aldrich #A1595). After 5 minutes at room temperature, cell viability was assessed by DAPI (Wako, #340-07971)/propidium iodide (PI, nacalai tesque, #19174-31) staining. Trypsin was used because, in preliminary trials, dissociation by pipetting in CMF alone required repeated mechanical agitation, which increased the number of PI-positive cells.

Dissociated cells were transferred to 15 mL low-binding conical tubes and fixed with 4.5 mL ACME solution (16.7% methanol, 11.1% acetic acid, 11.1% glycerol, 0.17% N-acetyl-L-cysteine/water) [29] for 20 minutes at room temperature with gentle rocking. Fixed cells were sequentially filtered through 40 µm and 20 µm cell strainers, then washed with 5 mL PBS containing 1% BSA and 0.2 U/µL RNase inhibitor (Protector RNase Inhibitor, Roche #3335399001). Cells were collected by centrifugation at 800×g for 5 minutes at 4°C, resuspended in 100 µL of 1% BSA/PBS with 0.2 U/µL RNase inhibitor, and pooled. The final cell suspension was filtered through a 40 µm strainer (Bel-Art Products, #H13680-0040) and adjusted to a concentration of 1,100–1,200 cells/µL. Cell concentration was confirmed by DAPI staining.

### Nuclei extraction for snRNA-seq

Ten adult *P. naikaiensis* individuals were collected on ice and washed twice with 500 µL ice-cold ASW, with 5-minute incubation on ice for each wash. Washed samples were transferred to 250 µL ice-cold lysis buffer (10 mM Tris-HCl pH 7.5, 10 mM NaCl, 3 mM MgCl₂, 0.1% IGEPAL CA-630 (Sigma-Aldrich #I8896), 1% BSA, 0.2 U/µL RNase inhibitor (Roche #3335399001)) and homogenised using a Power Masher II (Nippi #893002, #893061) with three cycles of 30-second homogenisation followed by 10-fold manual inversion.

Following homogenisation, 200 µL of the lysate was transferred to a fresh tube and subjected to vigorous pipetting for 5 minutes on ice to ensure complete nuclear release. Nuclei were pelleted by centrifugation at 500×g for 10 minutes at 4°C, and the supernatant was carefully removed. The pellet was resuspended in 500 µL nuclei wash buffer (1% BSA in PBS, 0.2 U/μL RNase Inhibitor) and pelleted again at 600×g for 10 minutes at 4°C. The final pellet was resuspended in 90 µL nuclei wash buffer, filtered through a 40 µm cell strainer (Bel-Art Products, #H13680-0040), and kept on ice. Nuclear concentration was adjusted to 1,100–1,200 cells/µL counted by DAPI staining.

### Library preparation, read alignment and quantification

Library preparations were conducted following 10x Genomics Chromium [Single Cell 3’ v3.1 Kit]. Sequencing was performed by the NGS core facility at the Research Institute for Microbial Diseases of the University of Osaka. The *P. naikaiensis* genome assembly and gene models were obtained from [30] (NCBI Accession: PRJDB7329). Two neural transcription-factor genes (*Otp* on scf2889, *Fez* on scf934) were added to the published gene models [30] with new gene IDs (g90001.t1, g90002.t1), and the annotation was converted from GFF to GTF with AGAT (v1.4.0). To improve 3′-UTR annotation for the 3′-based assay, gene models were extended with GeneExt [31]. Pooled cDNA (10x R2) reads were aligned to the genome with STAR [32] (--outFilterMultimapNmax 10), and the resulting alignments were used by GeneExt to extend 3′ ends of gene models. Of the 22,145 gene models, 7,269 were extended in scRNA-seq and 10,651 in snRNA-seq, with a median extension of 150 bp and 1,047 bp, respectively.

Reference packages were built with cellranger mkref (Cell Ranger v8.0.1; 10x Genomics) from the extended annotation. For scRNA-seq, we additionally built a reference combining the nuclear genome with the complete mitochondrial genome [30], both to test for a mitochondrial origin of the unmapped reads and to quantify mitochondrial content. The mitochondrial genome was annotated with MitoZ v3.6 using genetic code 9 (Echinoderm and Flatworm Mitochondrial), and the resulting annotation was merged with the nuclear annotation before 3′-UTR extension. Mitochondrial entries were reinstated after GeneExt processing, which had removed them. Reads were aligned and quantified per library with cellranger count (--expect-cells 10000, default parameters). Ambient and background RNA was removed from the raw count matrices with CellBender (remove-background; v0.3.2) [33]. For snRNA-seq, CellBender was run with default settings, in which the number of expected cells and total droplets included were estimated automatically from the data. For scRNA-seq, the automatically estimated cell number differed substantially from the Cell Ranger estimate, and these parameters were therefore set manually (--expected-cells 5000, --total-droplets-included 15000). Other parameters were left at their defaults (--fpr 0.01, --epochs 150) for both assays. Per-cell mitochondrial content (scRNA-seq) was computed in Seurat v5 [34] as the fraction of UMIs assigned to mitochondrial genes.

### Re-mapping of unmapped reads to *Tetraselmis* reference genomes

To investigate the origin of scRNA-seq reads that did not align to the *P. naikaiensis* reference, unmapped reads were extracted from the Cell Ranger BAM files with samtools v1.21 (view -f 4). Cell barcode (CB) and UMI (UB) tags were embedded into the read names to preserve per-cell information during alignment (paired-end suffixes removed to prevent STAR truncation), and reads ≥30 bp were retained. Two *Tetraselmis* reference genomes were used. *Tetraselmis striata* (NCBI: GCA_006384855.1, with JGI Phycocosm Tetstr1 gene annotation converted to GTF using AGAT v1.7.0) and *Tetraselmis* sp. (NCBI: GCA_057375155.1, genome-only). Reads were re-mapped with STAR v2.7.11b using parameters optimised for divergent-species alignment (--outFilterMismatchNoverReadLmax 0.06 --outFilterScoreMinOverLread 0.3). The mapping rate was calculated as the sum of uniquely and multi-mapped reads. Per-cell algal read counts were obtained by extracting cell barcodes from primary alignments (samtools view -F 256) [35] and added to the Seurat metadata.

### Quality control and doublet removal

Quality control was performed on the CellBender background-corrected matrices using ddqcR v0.1.0 [36], which applies adaptive, distribution-based thresholds (median absolute deviation) rather than fixed global cut-offs (initialQC followed by ddqc.metrics and filterData, with do.ribo = TRUE). Because mitochondrial genes are not annotated in the *P. naikaiensis* nuclear assembly, mitochondrial content was not used as a QC metric. Instead, the percentage of reads from ribosomal-protein genes (percent.rb) was included, based on a manually curated list of 44 ribosomal-protein genes. Putative doublets were then removed with DoubletFinder v2.0.4 [37]. After log-normalisation, scaling, and PCA, the optimal pK (0.005 for snRNA-seq and 0.29 for scRNA-seq, respectively) was selected by a parameter sweep (paramSweep and find.pK; pN = 0.25, PCs = 1:10), the expected doublet rate was set to 7.5% and adjusted for homotypic doublets using modelHomotypic on the cluster labels, and only cells classified as singlets were retained. The same procedure was applied to both libraries of each assay.

For scRNA-seq, an additional cluster-level filtering step was applied after the initial clustering. Although mitochondrial genes were not used for cell-level QC as described above, per-cell mitochondrial content was quantified using the nuclear + mitochondrial reference, and clusters with a median mitochondrial content exceeding 50% were considered to represent damaged or low-quality cells. Five of 20 initial clusters met this criterion and were removed (2,115 of 15,525 cells; 13.6%), and the remaining 13,410 cells were re-clustered (FindClusters, resolution = 0.8), yielding the 25 clusters used in all subsequent analyses.

### Normalisation, dimensionality reduction and clustering

The two libraries of each assay were combined with merge() (without integration or batch correction) and processed with the standard Seurat v5 workflow (v5.4.0) [34]: LogNormalise (scale factor 10,000), identification of 2,000 highly variable genes (FindVariableFeatures), scaling of all genes (ScaleData), and principal component analysis (RunPCA). The first 30 principal components were used for neighbour-graph construction (FindNeighbors), Louvain clustering (FindClusters, resolution = 0.8), and UMAP visualisation (RunUMAP). After clustering, the count layers were merged (JoinLayers). For scRNA-seq this yielded 13,410 cells in 25 clusters and for snRNA-seq, 18,702 nuclei in 22 clusters. Cluster markers were identified with FindAllMarkers (Wilcoxon rank-sum test, only.pos = TRUE), and genes with pct.1 > 0.2, adjusted *P* < 0.05 (Bonferroni), and avg_log_2_FC > 1 were retained as differentially expressed genes (DEGs).

### Silhouette score calculation

Silhouette scores were calculated independently for each method to assess clustering quality. For each dataset, Euclidean distances were computed from original PCA embeddings (30 principal components) obtained prior to integration using the silhouette function from the cluster R package (v2.1.4). Silhouette scores quantify how well each cell matches its assigned cluster compared to neighbouring clusters, ranging from -1 (incorrect assignment) through 0 (boundary between clusters) to +1 (well-separated cluster). Values > 0.5 indicate well-separated clusters. Statistical comparison between methods was performed using two-sided Wilcoxon rank-sum test (significance threshold: *P* < 0.05), with effect size quantified by Cohen’s *d* (|d| < 0.2: negligible; < 0.5: small; < 0.8: medium; ≥ 0.8: large).

### Gene selection

We manually curated 304 *P. naikaiensis* genes spanning major functional categories relevant to acoel biology (neuronal (*n* = 171) and non-neuronal (*n* = 133) genes). Gene selection was based on three complementary strategies, (i) reciprocal best BLAST hit (RBBH) searches against *Homo sapiens* (UniProt UP000005640), *Drosophila melanogaster* (UniProt UP000000803) reference proteomes using BLASTP (NCBI BLAST+ v2.14.0; E-value threshold 1e-3), (ii) cluster-based marker analysis on muscle (scRNA-seq clusters 9/11, snRNA-seq cluster 5) and ciliated cell clusters (scRNA-seq cluster 22, snRNA-seq cluster 14) with subsequent BLAST annotation filtering, and (iii) manual literature-based selection for genes with established functional roles in *P. naikaiensis* (e.g., PKD genes [38]). We additionally performed RBBH against the NCBI non-redundant protein database (nr) for this curated gene set to supplement functional annotation.

For RBBH searches, best hits in each species were then used as queries for reciprocal BLAST against the *P. naikaiensis* combined proteome (22,145 sequences: 22,143 from [30] plus two newly added *Otp* and *Fez*). Neuropeptide precursor genes (GnRH, Calcitonin) were identified using a targeted approach [39] rather than the proteome-wide BLAST search, because short, rapidly evolving neuropeptide precursors typically fail to produce significant BLAST hits. The list of curated genes, including BLAST statistics is provided in Table S2.

### Whole-proteome GO categorisation

To assess gene detection bias across the whole proteome, Gene Ontology (GO) annotations were transferred from *Homo sapiens* and *Drosophila melanogaster* reference proteomes to predicted *P. naikaiensis* proteins. All predicted protein isoforms of each *P. naikaiensis* gene were searched against the *H. sapiens* and *D. melanogaster* UniProt reference proteomes using DIAMOND v2.1.11 [40] in BLASTP mode with the --more-sensitive option (E-value cutoff 1 × 10⁻³, --max-target-seqs 5). The isoform with the lowest E-value across all isoforms of a gene was retained as the best hit. GO term annotations for the human and *Drosophila* best hits were obtained from the Gene Ontology Annotation files (goa_human.gaf and goa_fly.gaf) downloaded from the EBI GOA database [downloaded on 13/06/2026]. For each *P. naikaiensis* gene, the union of GO terms (Biological Process, Molecular Function, and Cellular Component) from its human and *Drosophila* best hits was assigned. Of the 22,143 predicted genes, 12,121 (54.7%) had a BLAST hit to either reference proteome.

Genes were then assigned a priori to eight functional categories (Neural, Muscle, Cilia, Mucosal/secretory, Digestion, Metabolism, Ribosome, DNA replication) based on 70 canonical, category-defining GO terms selected for each category (Table S4). Because many genes carry GO terms belonging to more than one category, all analyses were repeated using only genes assigned to a single category (exclusive-genes analysis; Fig. S5).

### Detection ratio comparison between methods

Detection rate analyses for both the curated 304-gene set and the whole-proteome GO categorisation followed an identical statistical framework. For each gene, the detection rate per method was defined as the fraction of cells/nuclei with non-zero expression in the raw count matrix. Method bias was quantified as the log_2_ ratio of detection rates, log_2_[(snRNA-seq% + 0.01) / (scRNA-seq% + 0.01)], where 0.01% was added to both detection ratios to avoid undefined values when a gene was undetected in one method. Differences in log_2_ ratio between neural and non-neural groups were assessed using Welch’s two-sample t-test and the Wilcoxon rank-sum test, with effect size quantified by Cohen’s d. Within each functional category, the mean log_2_ ratio was tested against zero with a two-sided one-sample t-test and confirmed with the Wilcoxon signed-rank test. *P*-values were corrected across categories using the Benjamini–Hochberg procedure. Analyses used the log-normalised RNA assay in Seurat v5.4.0 [34]. To assess robustness to multi-category assignments using GO categorised gene sets, all analyses were repeated restricted to genes uniquely assigned to a single GO category.

### Cluster-level correlation analysis

Highly variable genes (HVGs) were identified independently for each dataset with FindVariableFeatures function in Seurat using the vst method (top 2,000 features). To restrict the comparison to features that are reproducibly variable in both methods, we used the HVGs shared by both datasets (*n* = 671). For each shared HVG, mean expression was calculated per cluster in each dataset, and Pearson correlation coefficients were computed for all scRNA-seq and snRNA-seq cluster pairs. To test whether cluster size influenced correspondence, we correlated each snRNA-seq cluster’s cell count with its maximum (best-match) correlation (Pearson; *P* < 0.05).

### Identification of method-specific clusters

To identify cell populations recovered by only one method, we combined cross-method label transfer with CCA-based integration. First, bidirectional label transfer was performed between the scRNA-seq and snRNA-seq datasets (Seurat FindTransferAnchors and TransferData, 30 dimensions) [34]. For each transfer, the per-cell maximum prediction score was summarised as the median per original cluster. Prediction scores in the scRNA-seq→snRNA-seq direction were used to evaluate snRNA-seq clusters, and those in the snRNA-seq→scRNA-seq direction to evaluate scRNA-seq clusters. Clusters with a median prediction score < 0.7 were retained as candidates. Independently, the two datasets were integrated using Seurat’s anchor-based CCA integration (SelectIntegrationFeatures, FindIntegrationAnchors, and IntegrateData; 30 dimensions), and the integrated data were scaled, PCA-reduced, and clustered (FindClusters, resolution = 0.5). Each candidate was then evaluated against two criteria in the integrated space: (i) ≥80% of its cells fell within a single integrated cluster (concentration), and (ii) that integrated cluster was composed of ≥80% cells from the same method (method-dominance). Clusters meeting both criteria were defined as method-specific.

### Detection-ratio comparison of cluster markers

For each method-specific cluster, marker genes were identified with Seurat FindMarkers (Wilcoxon rank-sum test) and retained if expressed in >20% of the cluster’s cells (pct.1 > 0.2), enriched (avg log_2_FC > 1), and significant (adjusted p < 0.05, Bonferroni correction). For every marker, detection rates and the log_2_ ratio were computed as described above (see "Detection ratio comparison between methods"). For each cluster, the mean log_2_ ratio of its markers was tested against zero using a two-sided one-sample t-test, with *P*-values corrected across the six method-specific clusters using the Benjamini–Hochberg procedure.

### Identification of shared (reciprocal) clusters

To identify cell populations recovered concordantly by both methods, we applied a parallel four-criterion approach to the same label transfer and integration results described above. First, clusters with a median prediction score ≥ 0.7 were retained as candidates for correspondence. Each candidate was then evaluated against the same two integration-based criteria: (i) ≥80% of its cells fell within a single integrated cluster (concentration), and (ii) that integrated cluster showed balanced representation of both methods, defined as 20–80% composition by either method (mixing), following the thresholds used to classify integrated clusters (Fig. S7). As a third criterion, reciprocal correspondence was tested: for each candidate satisfying the above criteria, its main integrated cluster (the integrated cluster containing the largest fraction of its cells) was identified, and a snRNA-seq/scRNA-seq cluster pair was considered reciprocally matched if both clusters, identified independently within their own dataset, converged on the same main integrated cluster. As a fourth criterion, DEGs (defined as above: pct.1 > 0.2, adjusted *P* < 0.05, log_2_FC > 1) were compared between each reciprocally matched pair, and overlap was quantified using the Jaccard index (intersection size divided by union size). Pairs with a Jaccard index ≥ 0.10 were classified as strongly shared. Pairs meeting the first three criteria but falling below this threshold were classified as weakly supported.

## Results

### Quality assessment of scRNA-seq and snRNA-seq

We generated scRNA-seq and snRNA-seq datasets from whole *P. naikaiensis* adults (Fig. 1B–D). Against the *P. naikaiensis* nuclear reference genome, snRNA-seq reads mapped at 83%, while scRNA-seq achieved 50% mapping (Table S1). To test whether the unmapped scRNA-seq reads originated from mitochondrial transcripts, we remapped them to a reference supplemented with the mitochondrial genome. The mapping rate remained at 50% (Fig. 1C, Table S1), excluding a mitochondrial contribution. We retained the mitochondrion-supplemented reference for downstream scRNA-seq analyses to enable mitochondrial content as a QC metric, while snRNA-seq used the nuclear-only reference appropriate for nuclear isolation. Because *P. naikaiensis* established the symbiotic relationship with *Tetraselmis* algae (Fig. 1A) [28], we next hypothesised that the unmapped scRNA-seq reads might originate from co-captured algal RNA. To test this, we re-mapped the unmapped reads to *Tetraselmis* reference genome (Fig. S1A). As no genome is available for the *Tetraselmis* symbiont of *P. naikaiensis*, we used two publicly available *Tetraselmis* genomes: *T. striata* (GCA_006384855.1) and *Tetraselmis* sp. NKG400013 (GCA_057375155.1). Of the unmapped reads, 52% (Library 1) and 53% (Library 2) aligned to *T. striata*, and 36% and 37% aligned to *Tetraselmis* sp. NKG400013. The remaining unmapped reads likely reflect sequence divergence between the native symbiont and the available *Tetraselmis* references, as well as unannotated or non-coding sequences in the host genome. Per-cell algal read counts measured against the two references were strongly correlated (Pearson *r* = 0.995; Fig. S1B), indicating that the two *Tetraselmis* reference genomes yielded highly consistent estimates of relative algal RNA abundance across cells. Algal read abundance varied among host clusters (Fig. S1C–F). This may reflect the extracellular nature of this symbiosis [28] and/or ambient algal RNA released during the dissociation step, rather than indicating that all host cell types harbour intracellular symbionts. Re-mapping of the unmapped snRNA-seq reads to the *Tetraselmis* references also identified algal transcripts, although at levels more than six times lower than in scRNA-seq (Fig. S1A). This indicates that both dissociation methods co-capture symbiotic algal RNA, but to different extents.

After quality control, we retained 13,410 cells and 18,702 nuclei, grouped into 25 and 22 clusters, respectively (Fig. 1C, D). Both methods detected similar host gene numbers (20,143 and 21,565), but scRNA-seq detected more genes in >5% of cells (2,383 vs 1,443 genes), likely reflecting the capture of abundant cytoplasmic transcripts (Fig. 1C, Fig. S2). The per-cell gene detection was lower in scRNA-seq compared to snRNA-seq, while UMI counts were comparable between methods (median ∼370) (Fig. 1E).

To assess clustering quality independently for each method, we calculated silhouette scores from original PCA embeddings prior to integration. Silhouette score analyses showed that snRNA-seq produced better-defined clusters than scRNA-seq (mean: 0.175 vs 0.081; *P* < 2e-16; Cohen’s *d* = -0.38), with fewer negative scores (18.1% vs 37.1%) (Fig. S3). The lower clustering quality in scRNA-seq appears to partially result from the reduced sequencing depth for host transcripts caused by co-captured algal RNA (Fig. S1).

### Comparison of gene detection between two methods

To determine whether the two methods differ systematically in which genes they recover, we compared per-gene detection rates between scRNA-seq and snRNA-seq. Functional annotation in non-model invertebrates such as *P. naikaiensis* is often incomplete, with key gene classes (e.g., neuropeptide precursors) under-annotated by Gene Ontology (GO) transfer from distant model organisms. We therefore first manually curated 304 genes with well-established functions, identified by reciprocal BLAST searches, and grouped them into 11 functional categories: neurotransmitter synthesis, neuropeptide system, signalling proteins, neurotransmitter receptors, neurotransmitter transporters, transcription factors, mucosal/secretory-related, muscle, digestion (cathepsins), metabolism and cilia (Table S2; the number of genes in each category is shown in Fig. 2A). For each gene, we defined its detection ratio as the fraction of cells (or nuclei) expressing it, and quantified the method bias as the log_2_ detection ratios (snRNA-seq/scRNA-seq). Positive values indicate higher detection in snRNA-seq (Fig. 2A, Fig. S4, Table S3). Genes associated with neurotransmitter receptors showed higher detection ratios in snRNA-seq than in scRNA-seq, whereas metabolism-related and cilia-associated genes were more detected in scRNA-seq. Other neural genes with small sample sizes (neurotransmitter synthesis, neuropeptide system, and signalling proteins) showed similar trends of modest snRNA-seq enrichment, although individual category-level tests did not reach statistical significance. Grouping all 304 genes as neural (*n* = 171) or non-neural (*n* = 133) revealed a significant bias: neural genes were more enriched in snRNA-seq than non-neural genes (mean log_2_ ratio 0.75 vs -0.40; Welch’s t-test, *p* = 2.629e-07; Cohen’s *d* = 0.61) (Fig. 2B). Accordingly, 47% of neural genes versus 26% of non-neural genes showed >2-fold snRNA-enrichment, a trend driven largely by neurotransmitter receptors (Fig. 2B).

**Figure 2.**
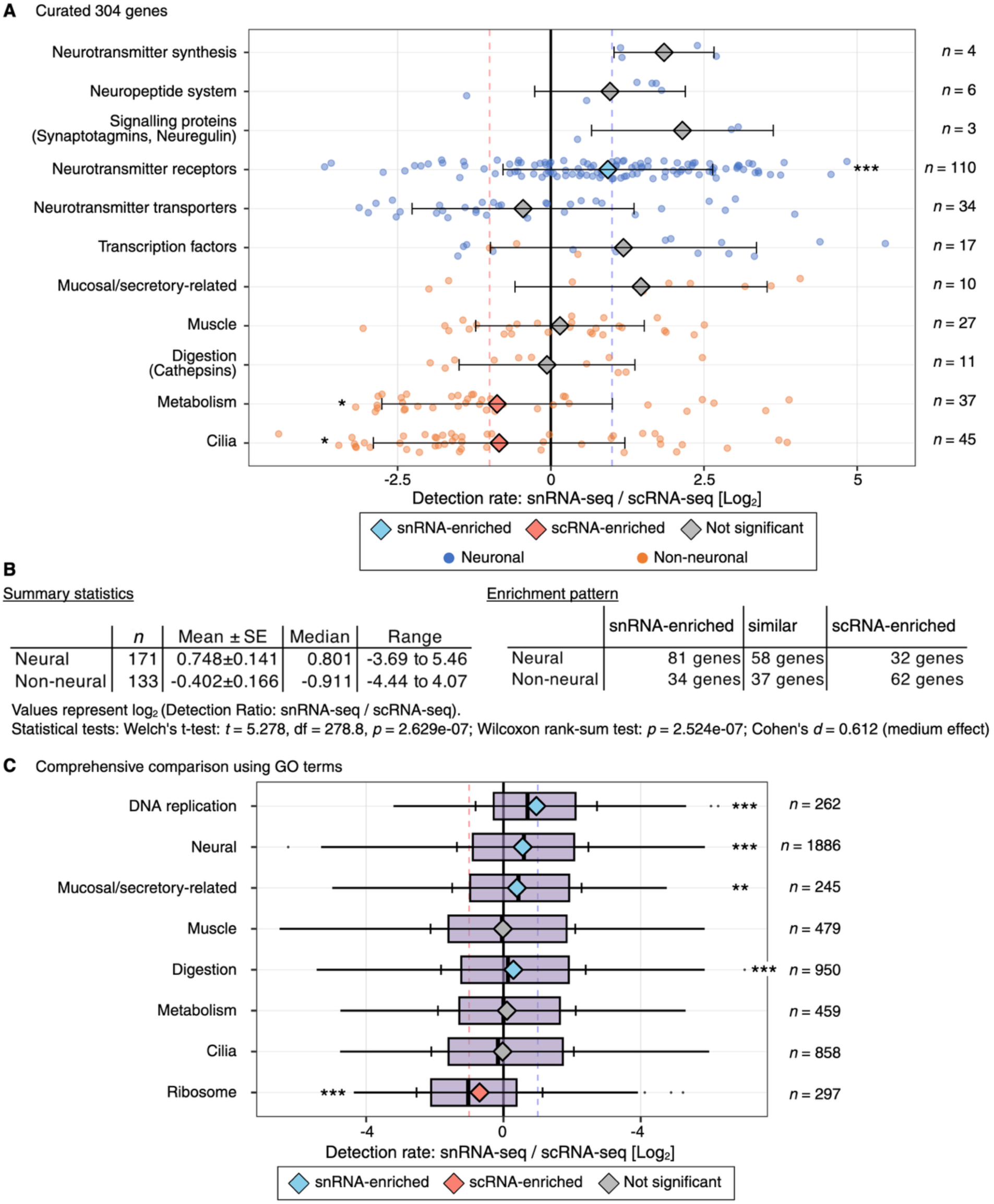
Comparison of detection levels of functional gene categories across snRNA-seq vs scRNA-seq. **(A)** Detection ratio comparison for 304 selected genes across 11 functional categories from snRNA-seq and scRNA-seq data in *P. naikaiensis*. Each point indicates a single gene, colored by neural (light blue) or non-neural (pink) classification. Diamond colors indicate light blue (snRNA-seq enrichment), pink (scRNA-seq enrichment), or gray (no significant difference). Dashed lines indicate 2-fold detection differences (log_2_ ratio = ±1). **(B)** Summary statistics comparing neural versus non-neural gene groups. Left table shows distribution statistics for log_2_ detection ratios. Right table shows enrichment patterns. Genes classified as snRNA-enriched (>2-fold), similar (within 2-fold), or scRNA-enriched (>2-fold). **(C)** Whole-proteome detection ratio by GO-defined functional category. Each *P. naikaiensis* gene with a BLAST hit to human or *Drosophila* reference proteomes was assigned a priori to one of eight GO categories (Table S4). Boxes indicate median and interquartile range. Diamonds indicate category means. Error bars indicate mean ± standard deviation. Diamond colors indicate category-level enrichment as in (A). Dashed lines indicate 2-fold detection differences (log_2_ ratio = ±1). Robustness analysis using genes uniquely assigned to a single category is shown in Supplementary Fig. 5. *** *P* < 0.001, ** *P* < 0.01, * *P* < 0.05, ns = not significant.

To test whether this enrichment is robust to the choice of curated genes, we extended the analysis to the whole proteome. GO terms were transferred to each predicted *P. naikaiensis* protein by DIAMOND searches against human and *Drosophila* UniProt proteomes, and genes were assigned a priori to eight functional categories based on canonical category-defining GO terms (Methods; Table S4). Neural genes (*n* = 1,886) showed significantly higher detection in snRNA-seq than scRNA-seq (Fig. 2C, Table S5), recapitulating the trend observed in the 304-gene set despite the broader, less stringent category definition. The neural enrichment was robust when the analysis was restricted to genes uniquely assigned to a single GO category (exclusive-genes analysis; Fig. S5, Table S5), confirming that inter-category overlap does not drive the observed pattern. The strongest snRNA-seq enrichment was observed for DNA replication-related genes. Notably, mucosal/secretory-related genes also showed significant snRNA-seq enrichment at the whole-proteome level. Similarly, the broader GO Digestion category (*n* = 950, encompassing diverse proteases and lysosomal hydrolases) was snRNA-seq enriched, whereas the cathepsin-focused curated digestion category (*n* = 11) showed no significant bias, suggesting that snRNA-seq enrichment characterises proteolytic genes more broadly than cathepsins alone. Conversely, ribosomal protein genes showed scRNA-seq enrichment, consistent with the predominantly cytoplasmic accumulation of ribosomal protein mRNAs [1]. This contrast highlights subcellular RNA localisation as a major determinant of the detection bias between the two methods. Pooling categories into neural (*n* = 1,886) versus non-neural (*n* = 2,802; genes uniquely assigned to one or more non-neural categories, excluding those also classified as neural) groups confirmed a significant overall bias toward snRNA-seq for neural genes (Welch’s t-test, *p* = 4.5e-12; Cohen’s *d* = 0.20). The smaller effect size compared to the curated 304 gene analysis (Cohen’s *d* = 0.61) reflects positive snRNA-seq enrichment in several non-neural categories at the whole-proteome level. Overall, snRNA-seq preferentially detects genes from functionally specialised cell populations, most prominently neural cells, but also secretory cells. In contrast, ribosomal transcripts, among the most abundant cytoplasmic RNAs, were significantly enriched in scRNA-seq (Fig. 2C), consistent with their cytoplasmic origin.

To assess cell-population correspondence between the methods, we correlated cluster-mean expression of the 671 highly variable genes (HVGs) across all scRNA-snRNA-seq cluster pairs and, for each snRNA-seq cluster, took its best-matching scRNA-seq cluster (Fig. S6). Best-match correlation ranged from *r* = 0.19 to 0.86: most of the snRNA-seq clusters (14/22) had at least a moderately correlated counterpart (*r* > 0.5), and this was independent of cluster size (*r* = 0.27, *p* = 0.22). Several snRNA-seq clusters shared the best-matching scRNA-seq cluster, suggesting that some populations resolved by snRNA-seq were merged into a single cluster in scRNA-seq. Thus, most snRNA-seq clusters (14/22) had a corresponding scRNA-seq population, while the remaining 8 clusters lacked a well-correlated counterpart (Fig. S6).

### A two-step approach identifies six snRNA-seq-specific clusters

To identify cell populations recovered by only one method, we combined cross-method label transfer with integration-based validation (Fig. 3A). Label transfer first flagged clusters that corresponded poorly to the other dataset, yielding 10 candidates (9 snRNA-seq, 1 scRNA-seq) (Fig. 3B). Because poor correspondence alone can also arise from rare or heterogeneous populations, candidates were additionally required to form a discrete, method-dominated group in the integrated data (Fig. 3A, Fig. S7, Table S6; see Methods for thresholds). This yielded six snRNA-seq-specific clusters (clusters 3, 4, 11, 16, 18, 19; 4,531 cells, 24.2% of snRNA-seq cells) and no scRNA-seq-specific cluster (Fig. 3C). Among the three excluded snRNA-seq candidates, clusters 1 and 2 dispersed across method-mixed integrated clusters and cluster 17 fell just below the concentration threshold. The single scRNA-seq candidate (cluster 22) concentrated in one integrated cluster, but that cluster was snRNA-dominant (71.5% snRNA) (Table S6, Fig. S7B).

**Figure 3.**
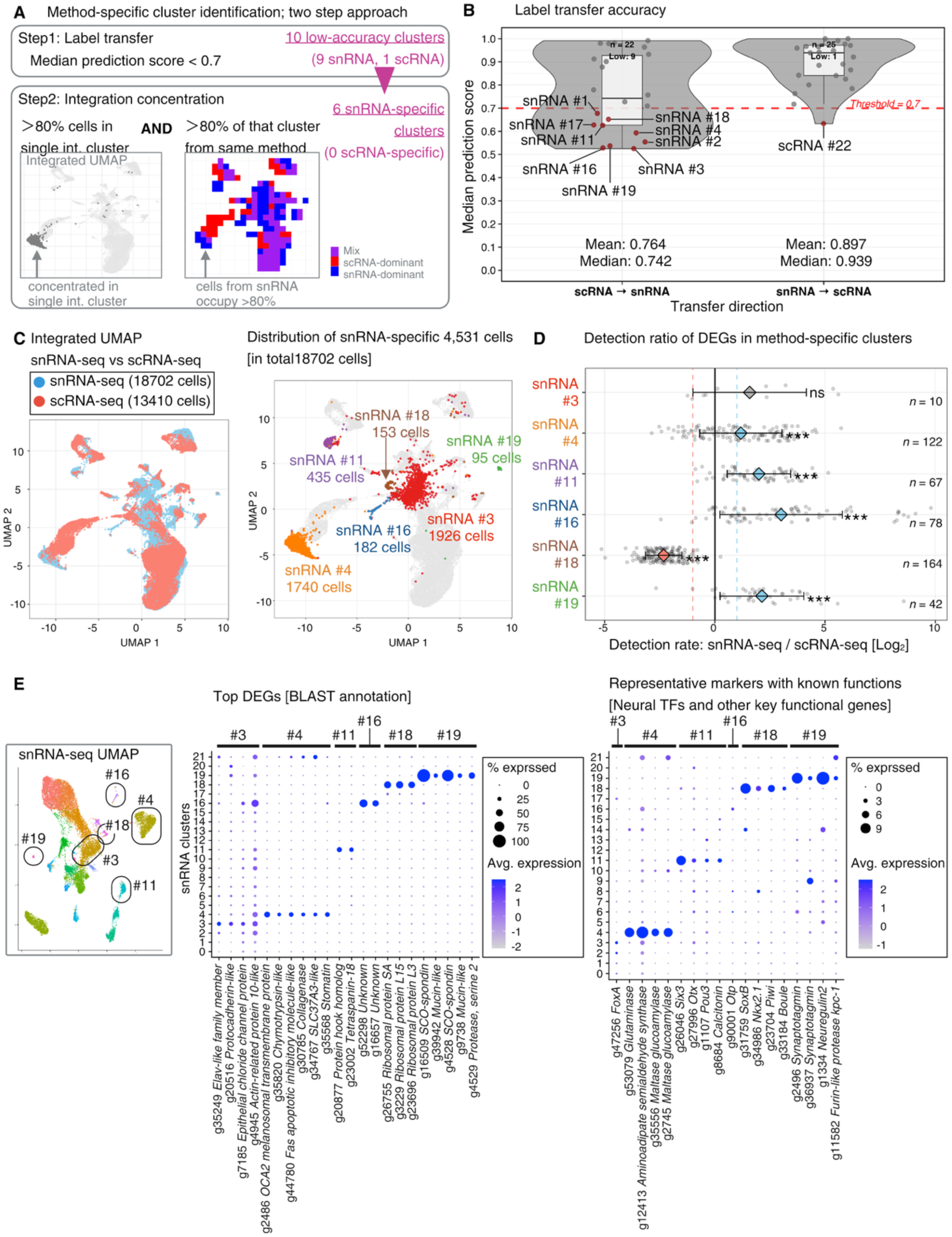
Identification and characterisation of method-specific clusters between scRNA-seq and snRNA-seq. **(A)** Two-step approach for identifying method-specific clusters. Clusters were first identified by low label transfer accuracy (median score < 0.7), then validated by concentration in integrated space (≥80% of cells in a single integrated cluster with ≥80% of that cluster from the same method). Numbers indicate clusters passing each criterion. **(B)** Label transfer accuracy. **(C)** Integration UMAP. **(D)** Detection of method-specific markers across methods. Log*_2_* detection ratio (snRNA-seq / scRNA-seq) for marker genes of the six method-specific clusters. Each point represents a qualified marker gene (pct.1 > 0.2, adjusted *P* < 0.05, log_2_FC > 1). Diamond colors indicate light blue (snRNA-seq enrichment), pink (scRNA-seq enrichment), or gray (no significant difference). Dashed lines indicate 2-fold enrichment thresholds (log_2_ ratio = ±1). *** *P* < 0.001, ns = not significant. **(E)** Gene expression of top differentially expressed genes with BLAST annotation (left panel) and manually annotated genes with known functions (right panel) across the six snRNA-specific clusters. Dot size represents the % of cells expressing each gene; dot colour indicates average scaled expression level. The two panels are shown separately because top DEGs and curated functional markers span markedly different ranges of expression frequency. Combining them on a single dot-size scale would render the low-frequency markers (e.g., TFs) invisible.

We next asked whether these clusters are method-specific because their defining genes are predominantly detected by one method. For each cluster’s DEGs, we computed the fraction of cells (or nuclei) of a given method in which each gene was detected and expressed the method bias as the log_2_ detection ratio (snRNA-seq/scRNA-seq) (Fig. 3D; DEGs and thresholds in Table S7 and Methods). Four clusters (4, 11, 16, 19) were significantly snRNA-seq-enriched, indicating that preferential snRNA-seq detection of their markers underlies their method-specificity, whereas cluster 3 showed no significant bias. Cluster 18 was unexpectedly scRNA-seq-enriched, its DEGs being dominated by abundantly and ubiquitously expressed ribosomal and housekeeping genes. Marker-level detection bias therefore does not explain its method-specificity, which instead arises at the level of its overall transcriptional profile (Fig. 3C).

### Molecular characterisation of method-specific clusters

To characterise the six snRNA-seq-specific clusters, we examined their marker genes using two complementary annotations. The top DEGs with BLAST-based annotations and a set of curated markers selected on biological grounds (Fig. 3E, left and right panels, respectively). Cluster 3 expressed the transcription factor (TF) *FoxA* together with an RNA-binding protein (CUGBP *Elav*-like), a protocadherin, and an actin-related cytoskeletal protein. Cluster 4 was enriched for *OCA2* (a melanosomal transmembrane protein), and a broad set of metabolic enzymes and transporters (*Chymotrypsin-2*, *Collagenase*, *SLC37A3*, *Stomatin*; and among curated markers, *Glutaminase*, *Aminoadipate-semialdehyde synthase*, and *Maltase-glucoamylase*), indicating a metabolically active population involved in amino-acid and carbohydrate metabolism. Cluster 11 expressed the neural TFs *Six3*, *Otx*, and *Pou3* and the peptide hormone *Calcitonin*, alongside structural and membrane proteins (*hook homolog*, *Tetraspanin-18*), consistent with a neural/neurosecretory identity. Cluster 16 had largely unannotated top DEGs but expressed the neural TF, *Orthopedia* (*Otp*). Cluster 18 was dominated by ribosomal-protein genes and co-expressed germline/stem-cell markers (*Piwi*, *Boule*) and the neural TFs, *SoxB* and *Nkx2.1*. This combination of stem markers (*Piwi*, *Boule*), neural developmental regulators (*SoxB*, *Nkx2.1*), and high ribosomal-gene expression is consistent with a proliferative progenitor-like population, potentially representing neural progenitors or an early neurogenic state. Cluster 19 expressed secreted and extracellular-matrix proteins (*SCO-spondin*, *Mucin*) and proteases (*Serine protease 2*, *furin-like protease kpc-1*) together with neural markers (*Synaptotagmin*, *Neuregulin*), consistent with neural or neurosecretory population.

Together, these six populations, neural, neurosecretory (clusters 11, 19), metabolic (cluster 4), and putative progenitor (cluster 18) identities, were detected exclusively or predominantly in snRNA-seq, indicating that nuclear profiling resolved cell types that are under-represented in our scRNA-seq data.

### Reciprocal cross-method correspondence identifies shared cell populations

Beyond the six snRNA-seq-specific clusters, snRNA-seq and scRNA-seq cells appeared well-mixed across most of the integrated UMAP (Fig. 3C, left panel), suggesting that many cell populations are concordantly captured by both methods. To identify these, we applied criteria parallel to, but inverted from, those above (Fig. 4A; see Methods). Clusters showing strong cross-method correspondence (Fig. 3B, Table S8) that concentrated within a single, method-balanced integrated cluster (Fig. S7). Applied independently to each dataset, these criteria identified six snRNA-seq and nine scRNA-seq clusters (Table S8). We then asked which of these converged reciprocally on the same integrated cluster from both directions, and finally required substantial DEG overlap between partners. Of four reciprocal pairs identified (1,933 scRNA-seq and 3,127 snRNA-seq cells; Table S9), three were retained: scRNA-seq cluster 9 / snRNA-seq cluster 5, scRNA-seq cluster 7 / snRNA-seq cluster 9, and scRNA-seq cluster 8 / snRNA-seq cluster 10. The fourth pair (scRNA-seq cluster 14 / snRNA-seq cluster 12) fell below the DEG overlap threshold and was excluded from further characterisation (Fig. 4B, Table S9).

**Figure 4.**
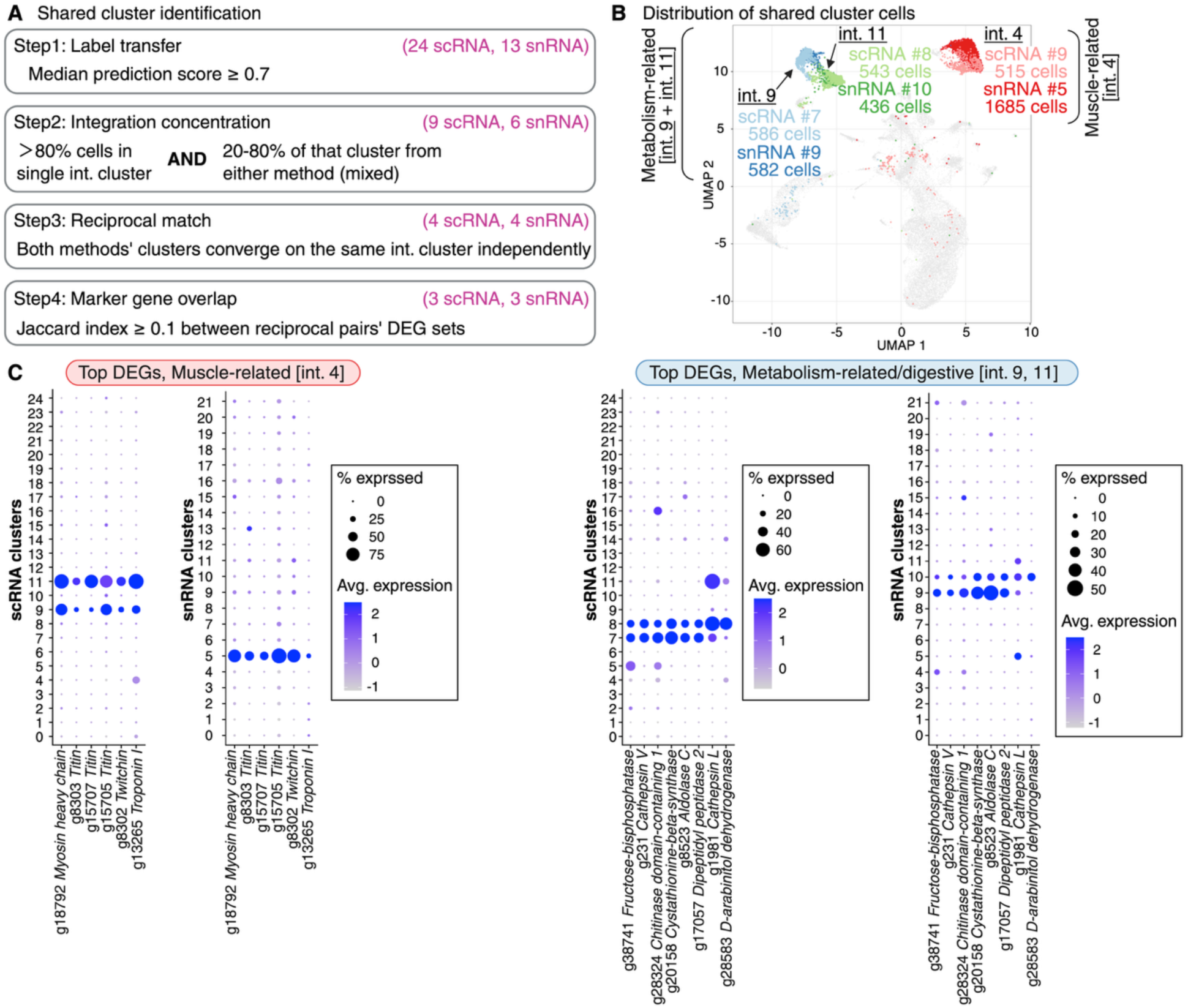
Identification and characterisation of shared clusters between scRNA-seq and snRNA-seq. **(A)** Four-step approach for identifying shared clusters. Numbers indicate clusters passing each criterion. **(B)** Integrated UMAP highlighting the three reciprocally matched shared cluster pairs, colored by pair identity and method of origin. Muscle: integrated cluster 4. Metabolic: integrated cluster 9 and 11 (see Fig. S6A for integrated cluster numbering). **(C)** Gene expression of top differentially expressed genes with BLAST annotation. Dot size represents the % of cells expressing each gene. Dot colour indicates average scaled expression level.

The three retained pairs shared coherent, functionally interpretable marker sets (Fig. 4C). The scRNA-seq cluster 9 / snRNA-seq cluster 5 pair shared core sarcomeric proteins (myosin heavy chain, titin, twitchin, and troponin I), consistent with a muscle cell identity independently recovered by both methods. Muscle-associated markers were also detected in scRNA-seq cluster 11, but this cluster did not satisfy the reciprocal correspondence criterion and was not considered part of the shared muscle population. The remaining two pairs shared several metabolic and proteolytic enzyme genes, including two cathepsins (cathepsin V and cathepsin L), fructose-bisphosphatase, cystathionine-beta-synthase, and dipeptidyl peptidase 2, suggesting closely related, metabolically active, potentially digestive cell populations. These genes were largely restricted to the four clusters comprising these pairs and were rarely detected elsewhere across the full dataset (Fig. 4B, C), arguing against non-specific, broadly expressed transcripts as the basis of this correspondence.

## Discussion

This study presents the technical comparison of scRNA-seq and snRNA-seq in a photosymbiotic invertebrate, revealing not only the method biases as observed in mammals, but also novel technical considerations for symbiotic systems.

Neural genes showed an overall tendency toward higher detection in snRNA-seq (Fig. 2A, B), consistent with observations from mammalian systems [1, 8–10]. Direct comparisons of scRNA-seq and snRNA-seq have also been performed in *Drosophila* [11, 12], but these studies did not specifically examine neural gene detection. Our results therefore provide, to our knowledge, the first evidence that this neural enrichment pattern extends beyond mammals. Significant enrichment was observed primarily for neurotransmitter receptor genes, whereas other neural categories, including neurotransmitter synthesis, neuropeptide-system-related genes and transcription factors, exhibited similar trends (Fig. 2A, S4). This suggests that snRNA-seq may capture transcripts of neural signalling genes more efficiently than scRNA-seq. In mammalian tissues, the advantage of snRNA-seq has been attributed partially to difficulties in dissociating tissue embedded in dense ECM [4, 6, 8]. However, acoelomorph nervous systems have minimal ECM [13, 19, 20], suggesting that neural enrichment in snRNA-seq cannot be explained solely by ECM-associated dissociation biases. The similarity of neural enrichment across phylogenetically and anatomically distinct systems suggests that snRNA-seq offers inherent advantages for neural cell profiling. Potentially, neural fragility during dissociation [41, 42], recapitulation of gene activity including immediate early genes [43], or differences in transcript localisation between nuclei and whole cells may contribute.

We also observed modest enrichment of cilia-associated and metabolic genes in scRNA-seq, potentially reflecting their cytoplasmic localisation.

The method-specific clusters were assigned from marker gene expression and their full cellular identities remain tentative (Fig. 3). While several clusters expressed marker combinations suggestive of neural or neurosecretory (snRNA-seq clusters 11, 19), metabolic (snRNA-seq cluster 4), or progenitor-like states (snRNA-seq cluster 18), definitive assignments will require future validation through in situ hybridisation. Nevertheless, the primary objective of this study was to evaluate how sequencing methodology influences cell type detection. From this view, six snRNA-seq clusters (Fig. 3C), representing ∼24% of all snRNA-seq profiles, were not resolved as distinct clusters in scRNA-seq. This is consistent with method-dependent differences in cell type composition reported in mammalian brain tissue[1–3, 10], kidney[6], and *Drosophila* eye-antennal discs[12]. Unlike other method-specific clusters, cluster 18’s marker genes were more detected in scRNA-seq than in snRNA-seq (See the dotplot in Fig. 3D). This indicates that its detection as a separate cluster in snRNA-seq is not explained by gene-level detection bias toward snRNA-seq, and instead may reflect a property of the cell population itself. One possibility is that this cluster represents progenitor-like transcriptional state characterised by co-expression of *SoxB*, *Nkx2.1*, *Boule* and one of the *Piwi* genes. *Piwi* has been reported as a neoblast marker expressed in pluripotent stem cells in the acoel *Hofstenia miamia* [44]. In another acoel *Symsagittifera roscoffensis*, neoblast-like cells localise along the nerve cords, and some are observed undergoing differentiation in the brain and epidermis [23]. If cluster 18 represents a progenitor-like population, its preferential recovery may reflect the ability of nuclear transcriptomes to resolve developmental cell states that are less captured in scRNA-seq.

Conversely, the identification of reciprocally matched clusters (Fig. 4) indicates that our approach for defining method-specific clusters was not systematically biased toward over-calling differences. Notably, the metabolically active shared clusters were independently subdivided into two closely related sub-populations by both methods (scRNA-seq clusters 7/8; snRNA-seq clusters 9/10), providing convergent evidence for a genuine biological distinction rather than a method-specific artifact. Notably, not all metabolically active populations were concordantly detected by both methods. The snRNA-seq-specific cluster 4 was also enriched for metabolic and transport-related genes (glutaminase, maltase-glucoamylase, associated with amino-acid and carbohydrate metabolism, Fig. 3).

The significant difference in mapping rates reflects co-capture of symbiotic algal RNA in scRNA-seq. This issue may influence experimental design such as sequencing depth and cell-sorting strategies. Remapping of unmapped reads demonstrated that algal transcripts accounted for approximately 26% of all scRNA-seq reads but only <4.2% of snRNA-seq reads (Fig. S1A). The lower proportion in snRNA-seq may partially reflect the exclusion of intact algal cells and cytoplasmic algal mRNA during nuclear isolation, whereas intact algal cells contribute their full transcriptome in scRNA-seq. Additionally, the two protocols differed in sample preparation. scRNA-seq samples were dissociated with trypsin and fixed with ACME, whereas snRNA-seq samples underwent unfixed nuclear isolation. These procedural differences may also have contributed to the difference in algal read proportion, although their relative contribution cannot be determined from the present data. Algal reads were detected across all host clusters (Fig. S1C– F), and their distribution among cell types was not analysed in detail. Given the extracellular symbiosis of the *P. naikaiensis*–*Tetraselmis*, this pattern may reflect broad distribution of *Tetraselmis* across host tissues [22]. It could also reflect ambient algal RNA released during sample processing. The biological significance of symbiont-derived transcripts likely depends on the mode of symbiosis. In extracellular *P. naikaiensis*–*Tetraselmis* association [22, 28], algal cells are co-isolated together with host tissue, which reduces the host transcript recovery. At the same time, their transcripts can reveal the molecular state of symbiotic algae as they undergo change following symbiosis establishment (e.g., loss of the cell wall, flagella and eyespot [45–47]). In intracellular symbioses, by contrast, host and symbiont transcripts can be co-captured within the same cell. FACS-based isolation has been combined with scRNA-seq to increase sampling efficiency for the symbiotic cell population [48]. Thus, whether symbiont-derived reads are viewed as a technical burden or as biologically informative signal depends on the biological question being addressed.

Notably, all other published single-cell datasets for xenacoelomorphs have used scRNA-seq [44, 49–51], making our snRNA-seq data particularly valuable for identifying cell populations underrepresented in the previous studies. Cross-species comparisons of such data should, however, account for method-specific biases to avoid mistaking technical artifacts for biological divergence. Building on these resources, we are currently developing a comprehensive cell atlas of *P. naikaiensis*, integrating these sc/snRNA-seq data with spatial and morphological characterisation, to be reported in a forthcoming study.

## Conclusion

The datasets generated here provide a benchmark for evaluating single-cell and single-nucleus transcriptomic approaches in invertebrates with a simple body plan. They may facilitate future efforts to integrate their cell atlases across studies and species, while also informing experimental design in invertebrates and symbiotic systems. Thus, our results suggest that combining scRNA-seq and snRNA-seq may provide complementary perspectives on cellular identity, physiological state, and host–symbiont associations, an approach applied in other invertebrates such as a spider species [52] and echinoderms [53].

## List of abbreviations

*scRNA-seq*: Single cell RNA-sequencing
*snRNA-seq*: Single nucleus RNA-sequencing
*DEG*: Differentially expressed gene
*TF*: Transcription factor
*ECM*: Extracellular matrix
*HVG*: Highly variable gene

## Declarations

### Availability of data and materials

scRNA-seq and snRNA-seq data have been deposited with links to BioProject accession number PRJDB42881 in the DDBJ database. Processed data (CellBender-corrected count matrices and Seurat objects) are available in the Genomic Expression Archive (GEA) under accession number E-GEAD-1303. All analysis code and gtf files used in this study are available at https://github.com/ryo-d-nakamura/P.naikaiensis_scRNA_snRNA_comparison.

### Ethics approval and consent to participate

Not applicable.

## Consent for publication

Not applicable.

## Authors’ contributions

RN designed the study and conducted all experiments and analyses. RN, MD, KK, TosuS and YJL performed the nucleus/cell isolation for library preparation. KK prepared libraries for scRNA-seq and snRNA-seq. YS provided laboratory resources and facilities for KK’s work. RN analysed the sequencing data in discussion with SA and MH. ToshiS. contributed the annotation of a peptide gene. RN wrote the original draft. TaS, MH, YJL, TosuS and MD revised and edited the manuscript. All authors approved the final manuscript.

## Acknowledgements

We acknowledge the NGS core facility at the Research Institute for Microbial Diseases of The University of Osaka for the sequencing. The authors used AI-based tools to edit and refine the manuscript text, and to assist in writing scripts used for data processing.

## Additional files

Additional file 1: Supplementary Figures 1-7

Additional file 2: Supplementary Tables 1-9 (data for Tables S2, S4, S7–S9 provided as separate .xlsx files, their legends are included in Additional file 2)

