## Supplementary Figures for "Method-dependent biases in cell type detection between single-cell and single-nucleus RNA sequencing in the photosymbiotic acoel *Praesagittifera naikaiensis*"

**Additional file 1**

Ryo Nakamura et al.

**This file contains: Supplementary Figures 1-7.**

**Supplementary figures**

Supplementary Fig. 1
Supplementary Fig. 2
Supplementary Fig. 3
Supplementary Fig. 4
Supplementary Fig. 5
Supplementary Fig. 6
Supplementary Fig. 7

**A** Mapping rate of scRNA-seq/snRNA-seq reads to host and algae genomes

| | Reads | Mapped reads | Mapping rate | (% of full library) |
|---|---|---|---|---|
| scRNA-seq Library 1 | | | | |
| Full → *P. naikaiensis* | 396,109,461 | 199,441,874 | 50.35% | |
| Unmapped → *T. striata* | 196,667,587 | 102,557,251 | 52.15% | (25.89%) |
| Unmapped → *Tetraselmis* sp. | 196,667,587 | 70,940,179 | 36.08% | (17.91%) |
| scRNA-seq Library 2 | | | | |
| Full → *P. naikaiensis* | 351,156,117 | 175,409,752 | 49.95% | |
| Unmapped → *T. striata* | 175,746,365 | 92,917,686 | 52.87% | (26.46%) |
| Unmapped → *Tetraselmis* sp. | 175,746,365 | 64,420,464 | 36.65% | (18.35%) |
| snRNA-seq Library 1 | | | | |
| Full → *P. naikaiensis* | 328,377,144 | 274,411,081 | 83.57% | |
| Unmapped → *T. striata* | 53,966,063 | 13,900,522 | 25.75% | (4.23%) |
| Unmapped → *Tetraselmis* sp. | 53,966,063 | 9,327,921 | 17.28% | (2.84%) |
| snRNA-seq Library 2 | | | | |
| Full → *P. naikaiensis* | 314,900,569 | 261,419,138 | 83.02% | |
| Unmapped → *T. striata* | 53,481,431 | 12,994,903 | 24.29% | (4.13%) |
| Unmapped → *Tetraselmis* sp. | 53,481,431 | 8,520,478 | 15.93% | (2.71%) |

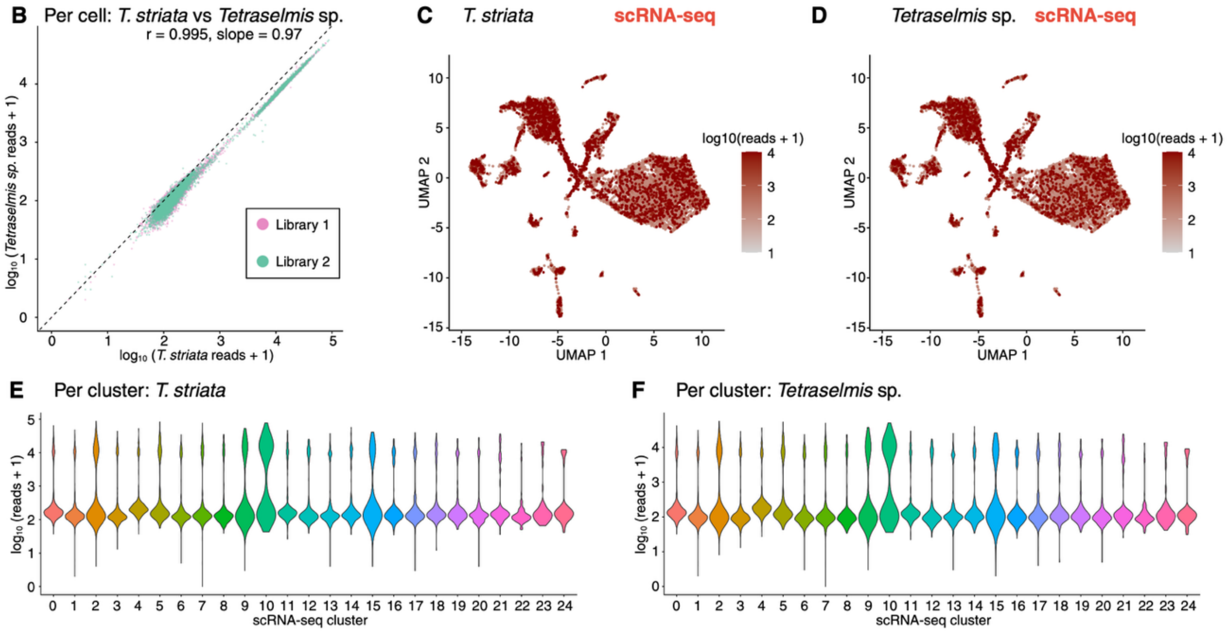


**Supplementary Fig. 1. Mapping of scRNA-seq unmapped reads to *Tetraselmis* algal reference genomes. (A)** Mapping rates of scRNA-seq/snRNA-seq unmapped reads to two *Tetraselmis* reference genomes. **(B)** Per-cell scatter plot of algal read counts mapped to *T. striata* (x-axis) versus *Tetraselmis sp*. (y-axis) in $\log_{10}$ scale (scRNA-seq). Points are colored by library identity. The dashed line indicates the diagonal (y = x). **(C, D)** UMAP visualization of host scRNA-seq cells ($n$ = 13,410), color-coded by the log10-transformed reaf count aligned to (C) *Tetraselmis striata* and (D) *Tetraselmis sp*. **(E, F)** Per-cluster distributions of algal reads ($\log_{10}$ scale) for (E) *T. striata* and (F) *Tetraselmis sp*. mapping in scRNA-seq.

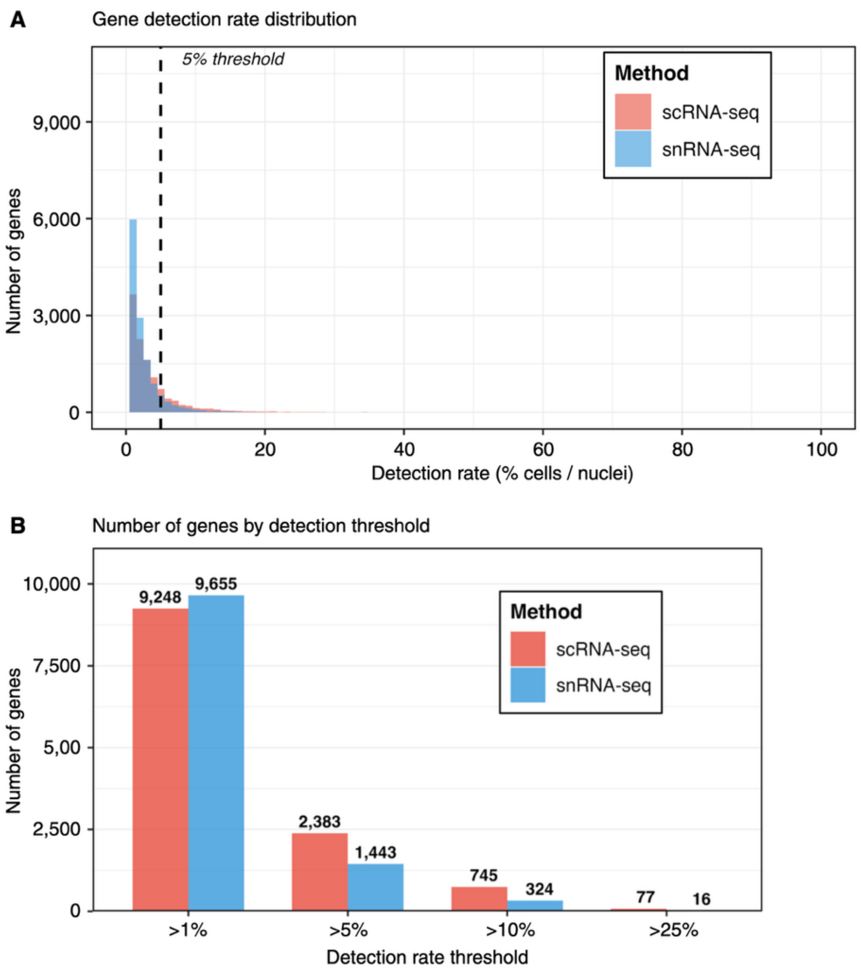


**Supplementary Fig. 2. Gene detection pattern from *P. naikaiensis* scRNA-seq and snRNA-seq data. (A)** Histogram showing the distribution of gene detection rates (% of cells or nuclei expressing each gene) for scRNA-seq (pink) and snRNA-seq (light blue). Dashed line indicates 5% detection threshold**. (B)** Number of genes detected above indicated thresholds. scRNA-seq detected more genes at the 5% threshold (2,383 vs 1,443 genes), reflecting enrichment for broadly expressed cytoplasmic transcripts. Numbers indicate gene counts at each threshold.

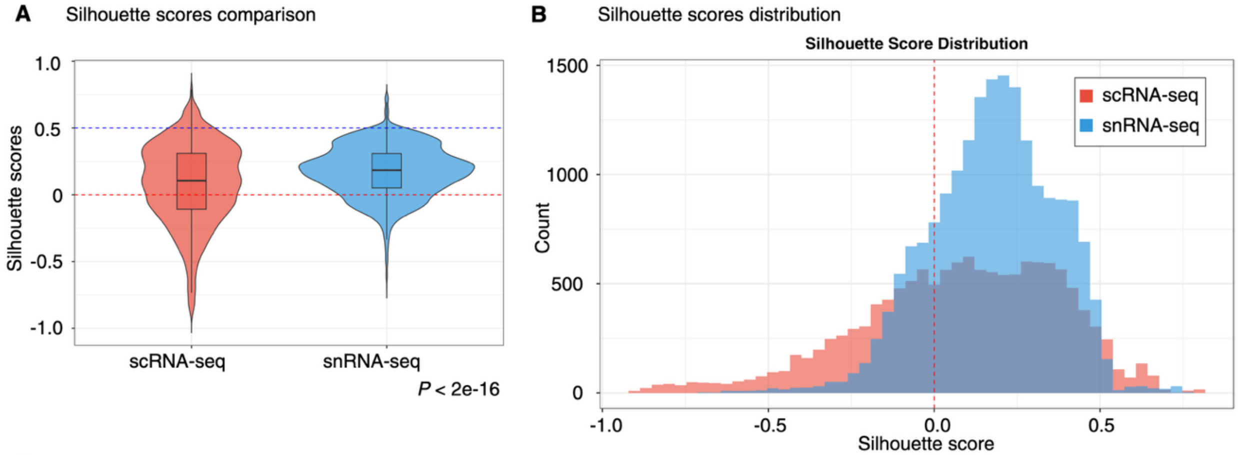


**C** Silhouette scores summary statistics

| Method | *n* (cells) | *n* (clusters) | Mean | Median | SD | Range | % Negative | % > 0.5 |
|---|---|---|---|---|---|---|---|---|
| scRNA-seq | 13410 | 25 | 0.081 | 0.106 | 0.298 | -0.913 to 0.791 | 37.10% | 5.4 |
| snRNA-seq | 18702 | 22 | 0.175 | 0.184 | 0.188 | -0.702 to 0.754 | 18.10% | 1.8 |

**Supplementary Fig. 3. Clustering quality comparison using silhouette scores. (A)** Violin plots showing silhouette score distributions for scRNA-seq and snRNA-seq. Box plots indicate median and interquartile range. Dashed lines: silhouette = 0 (red) and 0.5 (blue, well-separated clusters). **(B)** Histogram of silhouette score distributions. **(C)** Summary statistics. Silhouette scores were calculated from original PCA embeddings (30 dimensions) prior to integration. snRNA-seq showed significantly higher scores than scRNA-seq (Wilcoxon test, $P < 2\text{e-}16$; Cohen's $d = -0.38$, small effect).

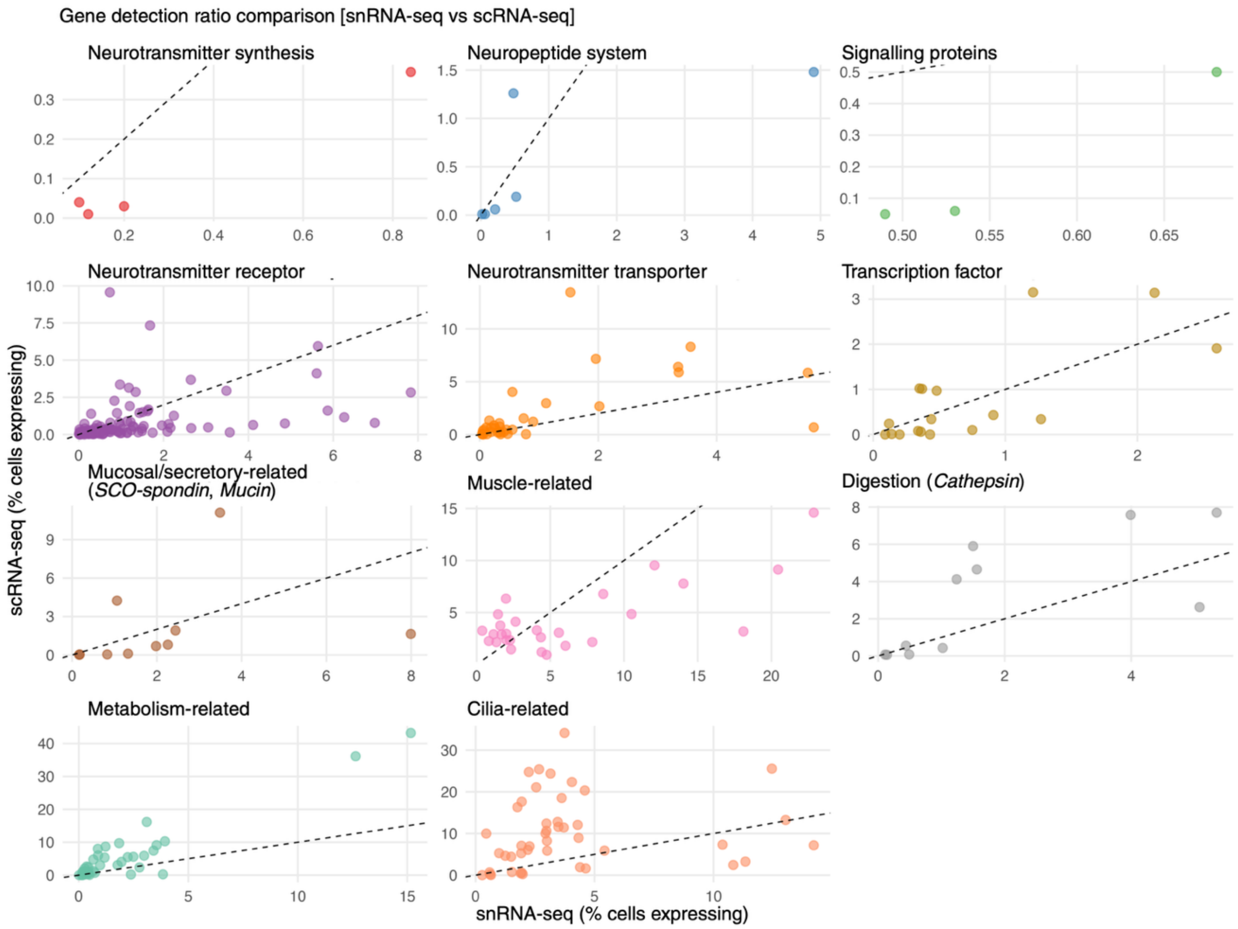


**Supplementary Fig. 4. Scatter plot comparison of gene detection ratios.** Scatter plots showing detection ratios (% of cells/nuclei expressing each gene) for 304 genes, comparing scRNA-seq (y-axis) versus snRNA-seq (x-axis) across functional categories. Each point represents one gene, colored by category. Dashed line: equal detection (y = x). Genes below the line show snRNA-seq enrichment. Genes above show scRNA-seq enrichment. Several categories show no systematic bias, with genes distributed around the diagonal. This gene-level view complements the category-level summary in Fig. 2A.

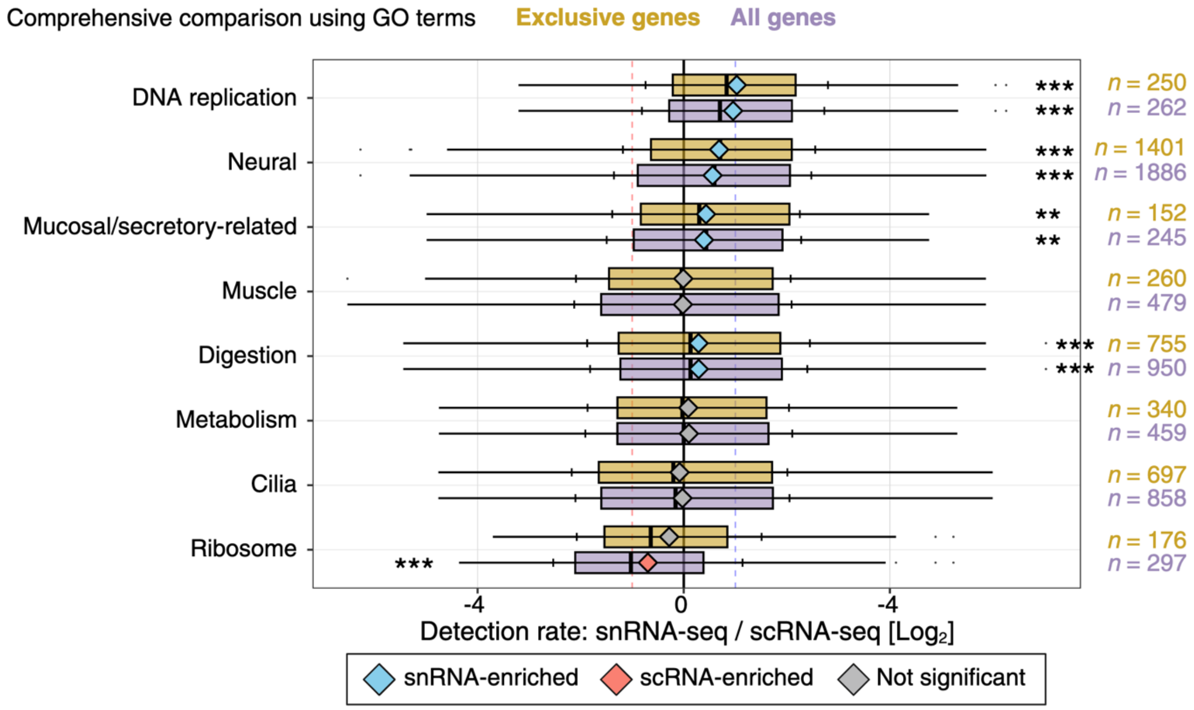


**Supplementary Fig. 5. Robustness of whole-proteome detection ratio comparison by GO category.** $Log_2$ detection ratios for genes assigned to eight GO functional categories, comparing "All genes" (multi-category assignments allowed, $n$ = 4,688 unique genes) with "Exclusive genes" (single-category only, $n$ = 4,031). *** $P < 0.001$, ** $P < 0.01$. Statistical elements as in Fig. 2C. Concordance between the two analyses confirms that inter-category overlap does not drive the observed patterns.

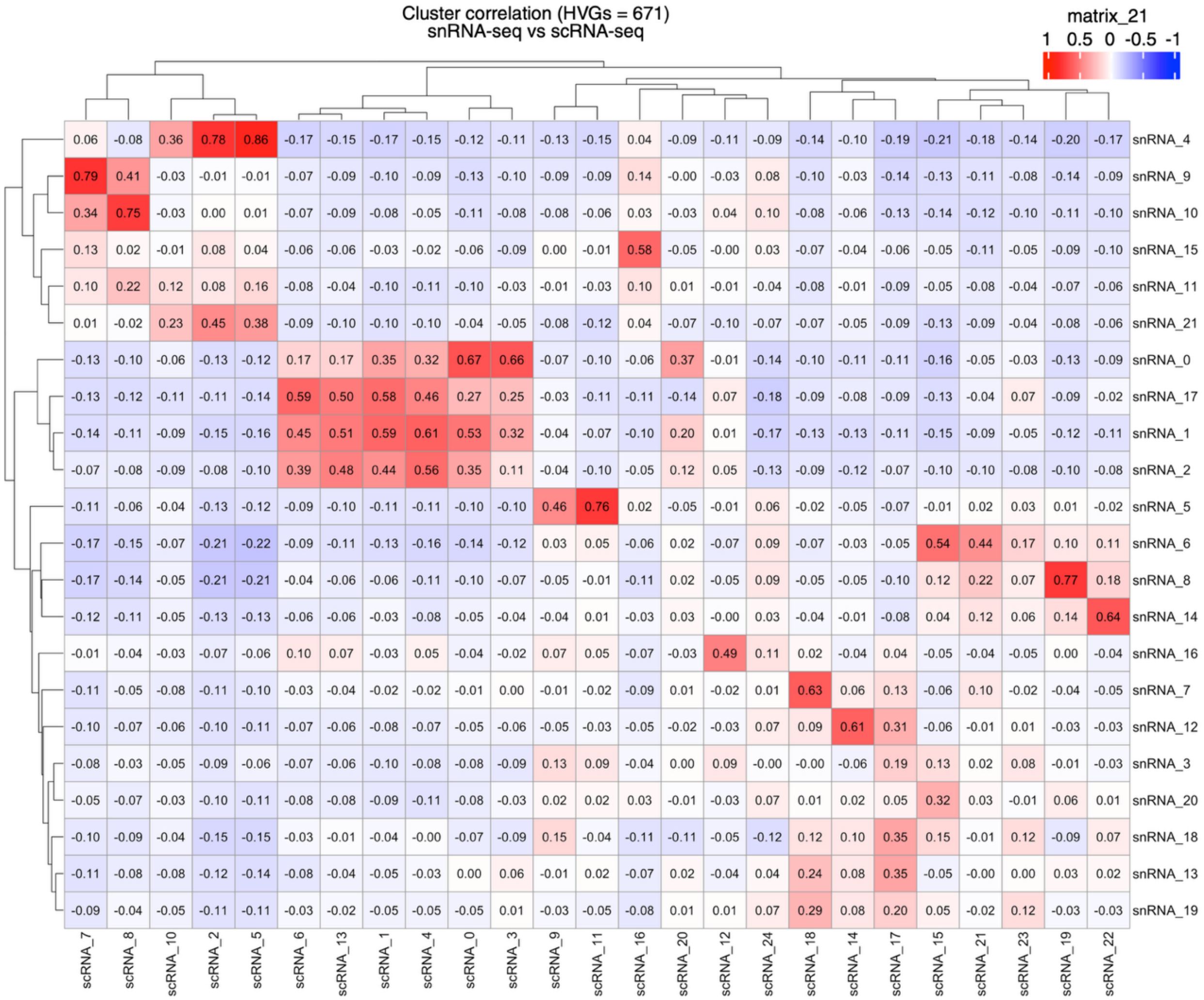


**Supplementary Fig. 6. Cluster-level correlation of common HVGs.** Heatmap showing Pearson correlation coefficients for cluster-level expression of HVGs detected in both scRNA-seq and snRNA-seq ($n$ = 671 genes). Each cell represents the correlation between one scRNA-seq cluster (columns) and one snRNA-seq cluster (rows). Colors indicate correlation strength: red (positive), blue (negative). Hierarchical clustering based on correlation patterns is shown on axes. Best- match correlations for each snRNA-seq cluster, showing wide variation ($r$ = 0.19 to 0.86). Correlation strength is independent of cluster size ($r$ = 0.27, $p$ = 0.22).

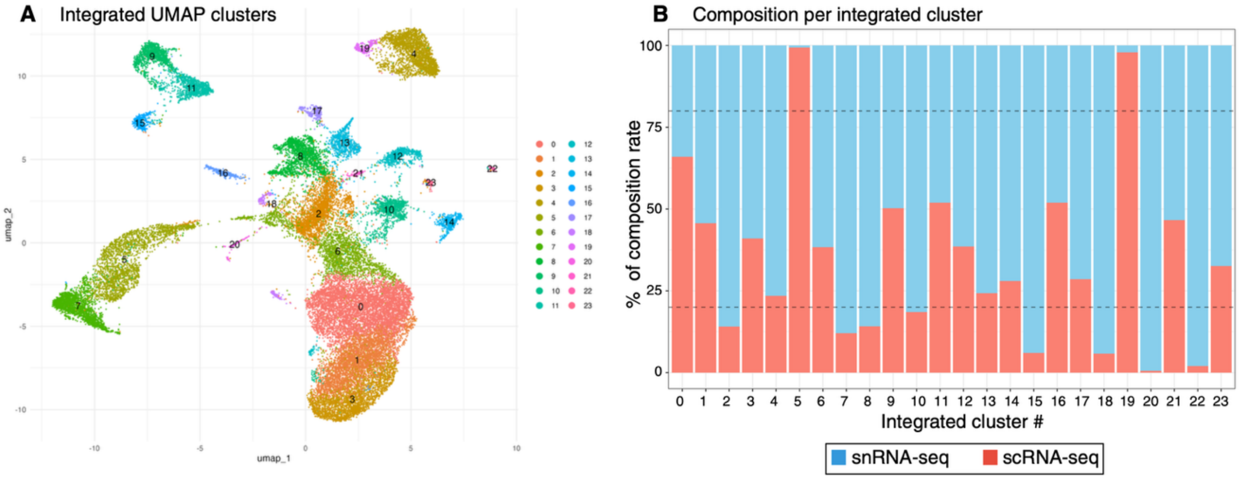


**Supplementary Fig. 7. Integrated UMAP clustering and dataset composition. (A)** UMAP of integrated scRNA-seq and snRNA-seq data, colored by integrated cluster number (0–23). Both methods were jointly clustered after integration ($n$ = 24 clusters). **(B)** Dataset composition per integrated cluster. Bars show % of scRNA-seq (salmon) and snRNA-seq (blue) cells in each cluster. Dashed horizontal lines indicate 20% and 80% thresholds used to define method-dominant clusters. Of 24 integrated clusters, 8 clusters (33.3%) were snRNA-seq-dominant (>80%), 2 (8.3%) were scRNA-seq-dominant, and 14 (58.3%) were mixed. Used to define method-specific spatial concentration in Fig. 3A.
