## Supplementary Tables for "Method-dependent biases in cell type detection between single-cell and single-nucleus RNA sequencing in the photosymbiotic acoel *Praesagittifera naikaiensis*"

**Additional file 2**

Ryo Nakamura et al.

**This file contains: Supplementary Tables 1, 3, 5, 6 (full tables) and legends of Supplementary Tables 2, 4, 7-9.**

**Supplementary tables**

Supplementary Table 1
Supplementary Table 2 (data provided separately as .xlsx; legend below)
Supplementary Table 3
Supplementary Table 4 (data provided separately as .xlsx; legend below)
Supplementary Table 5
Supplementary Table 6
Supplementary Table 7 (data provided separately as .xlsx; legend below)
Supplementary Table 8 (data provided separately as .xlsx; legend below)
Supplementary Table 9 (data provided separately as .xlsx; legend below)

| | Reference genome | | |
|---|---|---|---|
| | Nuclear only (%) | Nuclear + MT (%) | Difference (%) |
| scRNA-seq library 1 | 50.3 | 50.4 | +0.1 |
| scRNA-seq library 2 | 49.9 | 50.0 | +0.1 |

**Supplementary Table 1. scRNA-seq mapping rates with and without mitochondrial reference.** Mapping rate represents the % of reads mapped to the *P. naikaiensis* nuclear reference genome (using Cell Ranger). Adding the mitochondrial (MT) genome to the regerence resulted in a minimal change (+0.1%), indicating that the unmapped reads do not originate from mitochondrial regions.

**Supplementary Table 2. BLAST-based validation of 304 curated genes.** RBBH validation results for the 304 curated genes against *Homo sapiens* (Hsap) and *Drosophila melanogaster* (Dmel) reference proteomes, and the NCBI non-redundant (NR) protein database. For each gene, along with the best BLAST hit, E-value, bit score, query coverage (qcov), and percent identity (pident) for each database. NR symbol and description indicate the top NR database hit and its annotated identity, with corresponding accession number, E-value, bit score, and percent identity. *prepared as a separated excel file (Table_S2.xlsx)*

| Category | *n* | Mean $log_2$ | Median $log_2$ | t-test *p* | Wilcoxon *p* | t-test *p* (BH) | Wilcoxon (BH) | Significance (T-test *p*) *1 | Interpretation |
|---|---|---|---|---|---|---|---|---|---|
| Neurotransmitter synthesis | 4 | 1.847931 | 1.77689042 | 0.020157 | 0.125 | 0.05543158 | 0.171875 | ns | No difference |
| Neuropeptide system | 6 | 0.966137 | 1.52903453 | 0.112302 | 0.09375 | 0.17647487 | 0.147321429 | ns | No difference |
| Signalling proteins | 3 | 2.147508 | 2.94753258 | 0.12895 | 0.25 | 0.17730624 | 0.305555556 | ns | No difference |
| Neurotransmitter receptors | 110 | 0.931306 | 1.01023205 | 1.00E-07 | 2.81E-07 | 1.10E-06 | 3.09E-06 | *** | snRNA-enriched |
| Neurotransmitter transporters | 34 | -0.452678 | -0.82798163 | 0.154427 | 0.088917765 | 0.18874396 | 0.147321429 | ns | No difference |
| Transcription factors | 17 | 1.185624 | 1.06413034 | 0.038783 | 0.056884766 | 0.08532261 | 0.147321429 | ns | No difference |
| Mucosal/secretory-related | 10 | 1.472783 | 1.71667051 | 0.049881 | 0.083984375 | 0.09144895 | 0.147321429 | ns | No difference |
| Muscle | 27 | 0.149752 | 0.34055928 | 0.576527 | 0.49961102 | 0.63418004 | 0.549572122 | ns | No difference |
| Digestion | 11 | -0.062581 | -0.31550183 | 0.887842 | 0.965820313 | 0.8878421 | 0.965820313 | ns | No difference |
| Metabolism | 37 | -0.876467 | -1.29067716 | 0.007486 | 0.009616299 | 0.03111928 | 0.035259762 | * | scRNA-enriched |
| Cilia | 45 | -0.843384 | -1.49021189 | 0.008487 | 0.008814383 | 0.03111928 | 0.035259762 | * | scRNA-enriched |

**Supplementary Table 3. Per-category detection ratio test results of 304 curated genes.** Statistical results of the 304 curated gene set. Mean $log_2$ and Median $log_2$ represent the $log_2$-transformed fold-change in detection ratio between scRNA-seq and snRNA-seq datasets. *P*-values were calculated using a two-sided one-sample t-test against zero and a two-sided Wilcoxon signed-rank test against zero, with Benjamini-Hochberg (BH) correction applied for multiple comparisons across categories. *** $P < 0.001$, ** $P < 0.01$, * $P < 0.05$, ns = not significant. The 'Interpretation' column indicates whether each category shows detection enrichment in scRNA-seq, snRNA-seq, or no significant difference between the two methods.

**Supplementary Table 4. GO term definitions for whole-proteome analysis.** List of GO terms used in the whole-proteome analysis, categorised into eight functional categories (Neural, Muscle, Cilia, Mucosal_secretory, Digestion, Metabolism, Ribosome, and DNA_replication). For each GO term, the GO ID, GO term name, and GO aspect (Molecular Function, Biological Process, or Cellular Component) are provided. A total of 70 GO terms across 8 functional categories were used.

*prepared as a separated excel file (Table_S4.xlsx)*

**Exclusive genes**

| Category | *n* | Mean $log_2$ | Median $log_2$ | t-test *p* | Wilcoxon *p* | t-test *p* (BH) | Wilcoxon (BH) | Significance (T-test *p*) *1 | Interpretation |
|---|---|---|---|---|---|---|---|---|---|
| DNA replication | 250 | 1.028575 | 0.83138041 | 1.57E-17 | 2.67E-15 | 6.30E-17 | 1.07E-14 | *** | snRNA-enriched |
| Neural | 1401 | 0.685628 | 0.70866182 | 1.77E-40 | 4.87E-38 | 1.41E-39 | 3.89E-37 | *** | snRNA-enriched |
| Mucosal/secretory-related | 152 | 0.431983 | 0.29961576 | 0.003957 | 0.00818817 | 0.00791308 | 0.01310107 | ** | snRNA-enriched |
| Muscle | 260 | -0.008847 | -0.039995 | 0.945428 | 0.90914937 | 0.94542824 | 0.90914937 | ns | No difference |
| Digestion | 755 | 0.286595 | 0.12994641 | 0.000285 | 0.00085159 | 0.00076017 | 0.00227092 | *** | snRNA-enriched |
| Metabolism | 340 | 0.087779 | -0.0253205 | 0.408149 | 0.58258053 | 0.46645641 | 0.66580631 | ns | No difference |
| Cilia | 697 | -0.08167 | -0.2031487 | 0.303006 | 0.26916628 | 0.40400779 | 0.35888838 | ns | No difference |
| Ribosome | 176 | -0.282889 | -0.6402479 | 0.037768 | 0.00583015 | 0.06042938 | 0.0116603 | ns | No difference |

**All genes**

| Category | *n* | Mean $log_2$ | Median $log_2$ | t-test *p* | Wilcoxon *p* | t-test *p* (BH) | Wilcoxon (BH) | Significance (T-test *p*) *1 | Interpretation |
|---|---|---|---|---|---|---|---|---|---|
| DNA replication | 262 | 0.957378 | 0.70122107 | 2.54E-16 | 4.40E-14 | 1.02E-15 | 1.76E-13 | *** | snRNA-enriched |
| Neural | 1886 | 0.559477 | 0.59502215 | 1.89E-35 | 3.09E-33 | 1.51E-34 | 2.47E-32 | *** | snRNA-enriched |
| Mucosal/secretory-related | 245 | 0.391773 | 0.43521211 | 0.00131 | 0.002140439 | 0.00209601 | 0.003424703 | ** | snRNA-enriched |
| Muscle | 479 | -0.015991 | -0.04393 | 0.868249 | 0.824774463 | 0.86824922 | 0.824774463 | ns | No difference |
| Digestion (lysosomal protease) | 950 | 0.289839 | 0.13089793 | 2.46E-05 | 0.000101104 | 4.93E-05 | 0.000202208 | *** | snRNA-enriched |
| Metabolism | 459 | 0.098394 | 0 | 0.294151 | 0.446807765 | 0.39220169 | 0.595743687 | ns | No difference |
| Cilia | 858 | -0.025133 | -0.15947213 | 0.722994 | 0.647815088 | 0.82627856 | 0.740360101 | ns | No difference |
| Ribosome | 297 | -0.696168 | -1.02518724 | 2.82E-10 | 8.87E-11 | 7.52E-10 | 2.37E-10 | *** | scRNA-enriched |

**Supplementary Table 5. Per-category detection ratio test results of whole-proteome GO analysis.** Statistical results of whole-proteome GO-based analysis. GO terms used in the whole-proteome analysis are listed in Supplementary Table 4. N indicates the number of genes assigned to each category. Mean $\log_2$ and Median $\log_2$ represent the $\log_2$-transformed fold-change in detection ratio between scRNA-seq and snRNA-seq datasets. *P*-values were calculated using a two-sided one-sample t-test against zero and a two-sided Wilcoxon signed-rank test against zero, with Benjamini-Hochberg (BH) correction applied for multiple comparisons across categories. *** $P < 0.001$, ** $P < 0.01$, * $P < 0.05$, ns = not significant. The 'Interpretation' column indicates whether each category shows detection enrichment in scRNA-seq, snRNA-seq, or no significant difference between the two methods.

| Method | Cluster | *n* cells | Median prediction score | Int. Cluster ID | % cells in main int. cluster | % cells from same method | Method specific | Exclusion reason | Biological features |
|---|---|---|---|---|---|---|---|---|---|
| snRNA-seq | snRNA cluster 3 | 1926 | 0.525 | Int_2 | 83.1 | 85.9 | YES | | Largest cluster |
| snRNA-seq | snRNA cluster 4 | 1740 | 0.594 | Int_7 | 98 | 88 | YES | | |
| snRNA-seq | snRNA cluster 11 | 435 | 0.626 | Int_15 | 97.5 | 94 | YES | | TFs: *Six3*, *Otx*, *POU3*; *Calcitonin* |
| snRNA-seq | snRNA cluster 16 | 182 | 0.529 | Int_20 | 95.6 | 99.4 | YES | | |
| snRNA-seq | snRNA cluster 18 | 153 | 0.652 | Int_18 | 82.4 | 94.2 | YES | | Ribosomal genes, TFs: *SoxB*, *Piwi* |
| snRNA-seq | snRNA cluster 19 | 95 | 0.537 | Int_22 | 98.9 | 98.9 | YES | | Neuronal: *Synaptotagmin*, *Neuregulin* |
| snRNA-seq | snRNA cluster 1 | 2497 | 0.678 | Int_0 | 53.5 | 34 | NO | Scattered *1 | |
| snRNA-seq | snRNA cluster 2 | 2039 | 0.554 | Int_6 | 55.1 | 61.6 | NO | Scattered *1 | |
| snRNA-seq | snRNA cluster 17 | 165 | 0.628 | Int_18 | 72.1 | 94.2 | NO | Insufficiently concentrated *2 | |
| scRNA-seq | scRNA cluster 22 | 93 | 0.633 | Int_17 | 93.5 | 28.5 | NO | Not dominant *3 | |

*1 Cluster did not meet concentration criterion (>80% in single integrated cluster)
*2 Cluster fell below the concentration threshold
*3 Integrated cluster dominated by cells from the opposite method

**Supplementary Table 6. Evaluation of method-specific clusters.** All clusters with median transfer score < 0.7 were evaluated for method-specific detection using two criteria: (i) low label transfer accuracy (median score < 0.7), and (ii) concentration in integrated (>80% of cells mapping to a single integrated cluster, with >80% of that integrated cluster coming from the same method). Six snRNA-seq clusters met both criteria and were classified as method-specific. Three snRNA-seq clusters showed low transfer accuracy but were scattered across multiple integrated clusters, suggesting they represent heterogeneous or rare cell populations rather than method-specific detection. One scRNA-seq cluster (cluster 22) showed low transfer accuracy, but the corresponding integrated cluster was dominated by snRNA-seq cells, indicating cross-method detection rather than scRNA-specific detection.

Columns:
- Median prediction score: Median prediction score from label transfer (scRNA-seq→snRNA-seq direction for snRNA-seq clusters; snRNA-seq→scRNA-seq direction for scRNA-seq clusters)
- Int. cluster ID: Main integrated cluster after Seurat integration
- % in Main int cluster: Percentage of original cluster mapping to the main integrated cluster
- % from Same method: Percentage of cells in the integrated cluster from the same method as the original cluster
- Method specific: Whether cluster passed both criteria
- Biological features: Notable marker genes or characteristics

**Supplementary Table 7. All DEGs identified in snRNA-seq and scRNA-seq clusters, with annotations.** DEGs were identified for each cluster in both datasets using Seurat's FindAllMarkers, with genes defined as significantly differentially expressed if pct.1 > 0.2, $\log_2$ fold-change > 1, and adjusted *P*-value < 0.05. For each DEG, the best-matching BLAST hit (lowest E-value, E ≤ 1e-3) identified via DIAMOND BLASTP searches against the human and *D. melanogaster* proteomes is provided.

*prepared as a separated excel file (Table_S7.xlsx)*

**Supplementary Table 8. Cross-method correspondence and integration statistics for all clusters.** Median transfer score, main integrated cluster assignment, and integration concentration/composition statistics are shown for all 22 snRNA-seq and 25 scRNA-seq clusters. This table provides the complete underlying dataset from which method-specific (Table S6) and shared/reciprocal (Table S9) clusters were identified using the criteria described in Methods.

Columns:

- Median prediction score: Median prediction score from label transfer
- Int. cluster ID: Main integrated cluster after Seurat integration (i.e., the integrated cluster containing the largest fraction of the original cluster's cells)
- % cells in main int. cluster: Percentage of the original cluster's cells falling within the main integrated cluster
- % cells from same method: Percentage of cells in that integrated cluster originating from the same method as the original cluster

*prepared as a separated excel file (Table_S8.xlsx)*

**Supplementary Table 9. Evaluation of reciprocally matched shared clusters**. All scRNA-seq and snRNA-seq clusters meeting the correspondence (median prediction score ≥ 0.7), concentration (≥80% in single integrated cluster), and mixing (20–80% composition, Fig. S7B) criteria were evaluated for reciprocal correspondence with the other dataset. Four pairs converged reciprocally on the same integrated cluster. Three pairs additionally shared a substantial number of overlapping DEGs (Jaccard index ≥ 0.10) and were classified as strongly shared populations (Pairs 1–3; Fig. 4C). One pair (Pair 4) met the reciprocal correspondence criterion but fell below the DEG overlap threshold and was excluded from further characterisation.

*prepared as a separated excel file (Table_S9.xlsx)*
